# Tn-Core: context-specific reconstruction of core metabolic models using Tn-seq data

**DOI:** 10.1101/221325

**Authors:** George C diCenzo, Alessio Mengoni, Marco Fondi

## Abstract

**Motivation:** Tn-seq (transposon mutagenesis and sequencing) and constraint-based metabolic modelling represent highly complementary approaches. They can be used to probe the core genetic and metabolic networks underlying a biological process, revealing invaluable information for synthetic biology engineering of microbial cell factories. However, while algorithms exist for integration of –omics data sets with metabolic models, no method has been explicitly developed for integration of Tn-seq data with metabolic reconstructions.

**Results:** We report the development of Tn-Core, a Matlab toolbox designed to generate gene-centric, context-specific core reconstructions consistent with experimental Tn-seq data. Extensions of this algorithm allow: i) the generation of context-specific functional models through integration of both Tn-seq and RNA-seq data; ii) to visualize redundancy in core metabolic processes; and iii) to assist in curation of *de novo* draft metabolic models. The utility of Tn-Core is demonstrated primarily using a *Sinorhizobium meliloti* model as a case study.

**Availability and implementation:** The software can be downloaded from https://github.com/diCenzo-GC/Tn-Core. All results presented in this work have been obtained with Tn-Core v. 1.0.

**Supplementary information:** Supplementary data are available at Bioinformatics online.

## INTRODUCTION

The chemical complexity of biological entities hampers a full understanding of life and, consequently, its characterization is one of the strongest motivations in systems biology. Constraint-based metabolic modelling (CBMM) [1] is a well-established tool to formally represent cellular metabolism at the genome-scale level (by means of Genome Scale Metabolic Reconstructions, GSMRs) and to derive reliable predictions [2]. Despite this approach having shown remarkable predictive capabilities over the years [3], there are constant efforts aimed at improving and customizing the procedures of CBMM analyses.

It is increasingly recognized that the complexity of modern GSMRs often masks their utility in various applications [4], and that most studies to date only focus on the core metabolic pathways of the organism [5, 6]. Furthermore, due to the scaling of computational complexity, many stoichiometric (e.g. elementary flux modes enumeration [7]) and/or dynamic approaches (e.g. kinetic modelling [8]) cannot be applied to GSMRs embedding thousands of reactions. As a result, algorithms have been implemented to reduce a GSMR to a core set of reactions necessary to produce a pre-defined phenotype(s) [4, 9–11]. These algorithms share a similar overall approach: they are reaction-centric and require a user-defined list of reactions, metabolites, and/or phenotypes that must remain in the core model. However, by not directly incorporating experimental data, the biological accuracy of these core models cannot be guaranteed.

As GSMRs generally incorporate as much of the cell’s metabolism as possible, regardless to the activity of the reaction in a given environment, additional constraints are required to accurately represent environment-specific metabolism. This can be accomplished by constraining GSMRs with –omics data sets. This most commonly involves integrating gene expression data, constraining the allowable flux across each reaction based on the expression level(s) of the corresponding gene(s) [12, 13]. Similarly, tools exist for combining GSMRs with proteomics [14], fluxomics [15], and metabolomics data [16]. Ultimately, these applications have a common goal: reducing a GSMR to a smaller model with only the reactions active in the specific condition.

High-throughput transposon mutagenesis and sequencing (Tn-seq) generates a genome-wide list of genes essential in a given environment [17]. Arguably, these data sets are the best experimental representation of which reactions are active in a given environmental condition. Combining core metabolic networks and Tn-seq can allow deep functional refinement of GSMRs to account for only those (core) reactions and genes active under the tested conditions.. From a synthetic biology viewpoint, the central metabolism of an organism is of paramount importance as it i) produces the precursors for all natural chemicals and ii) has a high capacity of pathway fluxes; as such, central metabolism can be exploited as a chassis for production of industrially important molecules [18, 19]. Consequently, a Tn-seq curated core metabolic model is of high value for synthetic biology attempts at engineering designing cell factories. Indeed, genome streamlining, i.e., the construction of cells with minimal genomes, is known to generate cells with improved biotechnological properties, including increased protein or metabolite production [20–24]. However, despite the highly complementary nature of Tn-seq and CBMM, we are unaware of a tool for generating context-specific models through the automated incorporation of Tn-seq data with GSMRs.

Here, we report the development of Tn-Core, a MATLAB toolbox for use with COBRA formatted metabolic models. Tn-Core is designed for the generation of gene-centric, context-specific core metabolic models consistent with experimental gene fitness data produced through Tn-seq experiments, or through both Tn-seq and RNA-seq data. Tn-Core can further be used to: i) evaluate potential redundancy in core metabolism (does not require Tn-seq data); ii) identify which of the alternate pathway(s) contributes to higher flux through the objective function; and iii) perform Tn-seq-guided refinement of the Gene-Protein-Reaction rules (GPRs) in a GSMR.

## IMPLEMENTATION

Tn-Core was developed to facilitate the generation of context-specific core metabolic models through the integration Tn-seq data, then expanded to further allow the integration of RNA-seq data and to examine core metabolic redundancy in the presence or absence of these data. The toolbox is written in Matlab and uses COBRA formatted models and the COBRA Toolbox [25]. Tn-Core is available as Supplementary Materials S1, and the current and future versions will be available through GitHub (https://github.com/diCenzo-GC/Tn-Core).The functionality of the entire toolbox has been validated on four machines, running three versions of Matlab (R2015b, R2016b, R2017a) and three distinct COBRA toolbox setups (openCOBRA downloaded between 12/2016 and 08/2017), suggesting that Tn-Core should work in a broad range of computing environments.

### Generation of core metabolic models

The pseudocode for Tn-Core is given in Algorithm 1, the main workflow is depicted in the flowchart of Figure 1, and a detailed manual describing its usage is provided in Supplementary Materials S1. The minimum input is a COBRA-formatted metabolic model. Optionally, the user may provide: (i) Tn-seq data for all genes in the genome; (ii) RNA-seq data for all genes in the genome, and/or (iii) a list of pre-determined core/essential genes. Tn-Core begins with the optional step (Figure 1a) of producing a list of model genes to be protected during the generation of random core models. This list is based on: (i) all user-defined core genes, (ii) highly expressed genes if RNA-seq are provided, and (iii) essential genes based on Tn-seq data (optional even if Tn-seq data are provided). Next, Tn-Core produces a reduced GSMR by iteratively removing all reactions that produce dead-end metabolites (and associated genes, if they are not in the GPR of another reaction). Additionally, all GPRs not assigned to a coding sequence (e.g. gap-filling reactions) are removed. As the order in which reactions are added/removed from a model might alter the predictive capability of the reconstruction, randomized core models (*M*, Algorithm 1) are then generated from the reduced model (Figure 1b). Importantly, this step can be parallelized, reducing the running time. This involves first preparing a list of all non-protected model genes (*U_m_*), and randomly shuffling their order at each iteration (*U_m_*\*). All genes (and corresponding reactions) from each shuffled set are individually deleted from the model and growth is tested. If the objective function flux (*φ*) stays above the threshold (*t*), the gene is excluded from the model; otherwise, the gene is put back to the model. The result is a population of models (*M*) each containing the initially protected genes (optional), and a minimal amount of additional genes required to maintain objective function flux *φ* above the threshold *t*. If Tn-seq data is provided, the objective function flux of each core model is recorded, genes are classified into four categories from ‘essential’ to ‘non-essential’ based on the Tn-seq data (Figure S1), and the number of core model genes in each category is recorded (Figure 1c).

**Figure 1.**
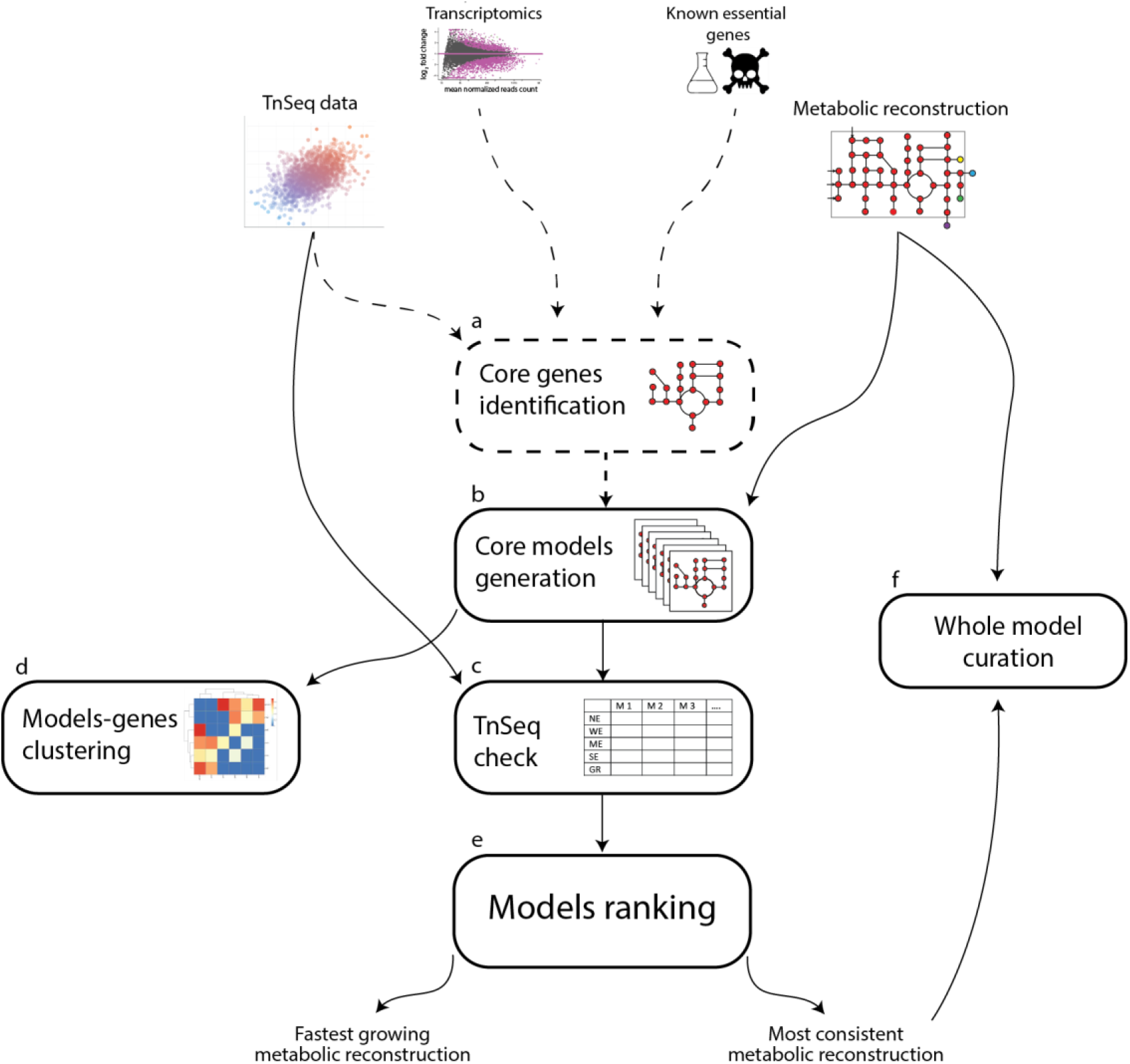
Schematic representation of the Tn-Core pipeline. Dashed lines represent optional steps.

#### Algorithm 1. The Tn-Core algorithm

**Input:** *n* is the number of iterations; *model* is the initial GSMR. **Other variables and lists:** *G_m_* is the list of genes in *model, T* is the Tn-seq data, *L* is the RNA-seq data, *M_i_* is the final array of core metabolic reconstructions, *t* is the threshold for objective function. **Functions:** *detectDeadEnds, deleteModelGene, findRxnsFromMets, singleGeneDeletion*, and *optimizeCbModel* are part of the COBRA Toolbox. All the other functions are implemented as Matlab code (see Supplementary Material S1).

~~~
**1:** *D* = ***detectDeadEnds***(*model*)
**2:** *R_D_* = ***findRxnsFromMets**(D)*
**3:** *m_red_* = **remove reactions and unused genes** (*model, R_D_*)
**4:** *E_model_* = ***singleGeneDeletion***(*model*)
**5:** (*E, S*, ***W***) = **get essential, strong, and weak growth promoting Genes** (*T*)
**6:** *L_H_* = **get highly expressed genes** (*L*)
**7:** *U_m_* = (*G_m_* ~ ((*E* ∩ *G_m_*) ∪ *E_model_* ∪ (*L_H_* ∩ *G_m_*)))
**8: for** *i* = 1 **to** *n*
**9:**                    *m* = *m_red_*
**10:**                   *U_m_*\* = shuffle(*U_m_*)
**11:                                  for** *j* = 1 **to** length(*U_m_*\*)
**12:**                                                  *m’* = *deleteModelGene*(*m, U_m_*\*(*j*))
**13:**                                                  *φ* = *optimizeCbModel*(*m’*)
**14:                                                                 if** φ > *t*
**15:**                                                                                *m* = *m’*
**16:                                                                 end if
17:                                        end for
18:**                         *M(i)* = *m*
**19:**                   *G_M(i)_* = **get the genes in** *M(i)*
**20:**                   *O(i)*= *optimizeCbModel(M(i))*
**21:**                      {*N_E_*(*i*); *N_S_*(*i*); *N_W_*(*i*)} = {**length**(*G_M(i)_* ∩ *E*); **length**(*G_M(i)_* ∩ *S*); **length**(*G_M(i)_* ∩ *W*)}
**22: end for
23:** *M_core_* = *M*(**max**(*N_E_*))
**24: if length**(*M_core_*) > *1*
**25:**    *M_core_* = *M_core_*(**max**(*N_S_*))
**26:**    *M_core_* = *M_core_*(**max**(*N_W_*))
**27:**    *M_core_* = *M_core_*(**max**(*O*))
**28: end if
29: return** *M_core_*
~~~

Finally, the core reconstruction that maximizes the number of essential Tn-seq genes is chosen as the reconstruction most consistent with the Tn-seq data (*M_core_*). If two or more models embed the same number of essential genes, the reconstruction maximizing the number of ‘strong growth promoting’ and then ‘weak growth promoting’ genes is selected as the output. If multiple models still remain, the model with the highest objective reaction flux is returned as the core metabolic model most consistent with the gene essentiality data (Figure 1d). Independently, the core model with the highest objective function flux is returned as the fastest growing core model (Figure 1d); if multiple models have the same maximal objective function flux, the model most consistent with the gene essentiality data is chosen. In some cases, it may be desirable to obtain other core models produced during the running of Tn-Core, such as the slowest growing core model. The output of Tn-Core additionally includes a cell array of the objective function flux for all produced core models, as well as a binary presence/absence cell array indicating which genes are included in each of the core models. By using the latter cell array with the *tncore_reconstruct* function, it is possible to rebuild any of the core models produced during the running of Tn-Core.

### Analysis of variation across the core metabolic models

The redundancy embedded within GSMRs means that each of the models in the core model population may contain a different set of genes and/or reactions. Tn-Core includes functions to explore this redundancy, whether Tn-seq data is provided or not (Figure 1e). Two or three primary matrixes are returned, and can display either gene or reaction information. A binary presence/absence matrix is given, which indicates, for each model, whether each feature is present or absent; only features embedded in at least one core model are included (Figure 2a, 2b). A co-occurrence matrix is also provided; for each feature variably present in the core model population, a Chi-squared statistics is reported to indicate which feature pairs are more likely than chance to appear, or not appear, in the same core models (Figure 2c-2e). If the core models are generated multiple times, for example, using different objective flux thresholds, a matrix can be produced that indicates, for each population of core models, what percentage of models contains each of the features (Figure 2f, 2g).

**Figure 2.**
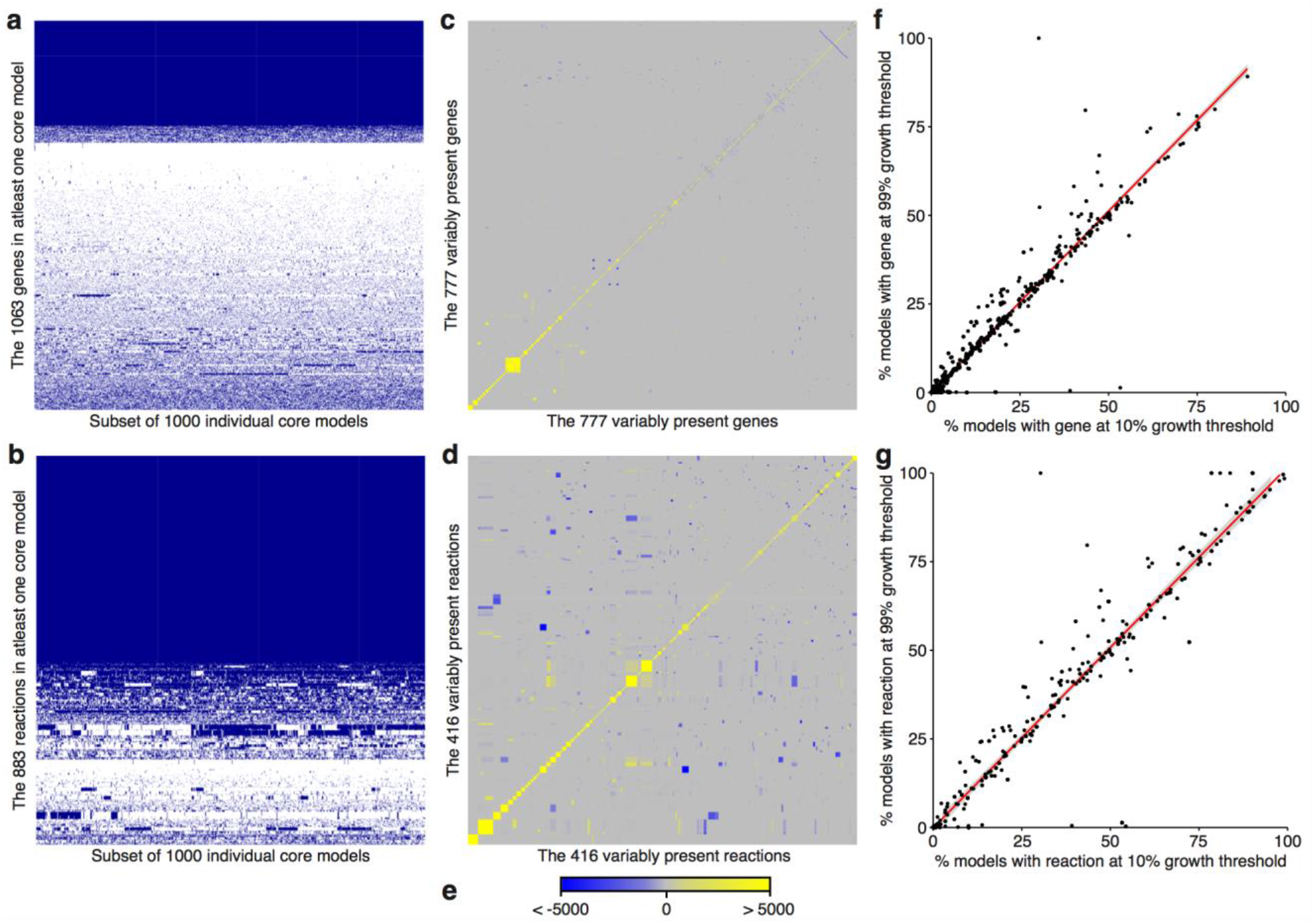
Evaluation of core metabolic redundancy with Tn-Core. The six primary matrixes generated by Tn-Core are shown. Tn-Core was run using the *S. meliloti* iGD1575 genome-scale metabolic reconstruction, with 25,000 iterations, a growth threshold of 10%, without essential gene pre-identified, and without RNA-seq data. Gene (**a**) and reaction (**b**) presence matrixes are shown for 1,000 of the randomly produced core models. Blue indicates the gene/reaction is present, white indicates the gene/reaction is absent. Gene (**c**) and reaction (**d**) co-occurrence matrixes are shown for the genes/reactions variably present in the 25,000 core models. (**e**) The legend for the co-occurrence matrixes is shown. The scale represent a Chi-squared statistic that summarizes if the gene or reaction pair is more (yellow) or less (blue) likely to occur in the same core model than by chance. Gene (**f**) and reaction (**g**) scatter plots displaying the correlation between the percentage of core models containing the gene/reaction when made using a growth threshold of 10% or 99%. Genes/reactions either present in all models or in no models are not included.

### Refinement of genome-scale metabolic network reconstructions

Finally, an extension is provided to use Tn-seq data to assist in the automated curation of GSMRs (Figure 1f). First, Tn-seq essential genes are determined, and these genes are protected during core model generation. The core model most consistent with the Tn-seq data is collected, and where appropriate, ‘or’ statements in the GPRs are replaced with ‘and’ statements; if any Tn-seq essential genes in the model have no effect when deleted, and if any occur in the same reaction(s) and only the same reaction(s), and the GPR currently lacks an ‘and’ statement, the ‘or’ statements of the GPR are replaced with ‘and’ statements. The implementation of this section of the code is rather strict in order to avoid artificially converting non-essential genes to essential genes. Finally, for any core model reaction with a Tn-seq essential gene, the corresponding GPRs of the original GSMR are replaced with those of the core reconstruction.

## RESULTS AND DISCUSSION

### Validation of Tn-Core

Tn-Core was validated by extracting context-specific core models from the *Sinorhizobium meliloti* iGD1575 GSMR [26]. Two core models were produced, each using a growth threshold of 50% the full model, with 50,000 iterations, and with Tn-seq essential genes pre-identified. In one Tn-Core run, only Tn-seq data [27] was used; in the second run, the same Tn-seq data plus RNA-seq data [28] was included. The sizes of both models are summarized in Table 1, and the inclusion of RNA-seq data resulted in a somewhat larger core model.

**Table 1.** Summary of the sizes of the produced core models relative to the parent model (iGD1575) and the manually prepared core model (iGD726).

| Model | Genes | Reactions | Metabolites |
| --- | --- | --- | --- |
| iGD1575 | 1577 | 1828 | 1579 |
| iGD726 | 728 | 681 | 703 * |
| Core model A (without RNA-seq, FBA) | 488 | 574 | 578 |
| Core model B (with RNA-seq, FBA) | 532 | 614 | 601 |
| Core model C (without RNA-seq, MOMA) | 490 | 581 | 584 |
| Core model D (with RNA-seq, MOMA) | 532 | 602 | 590 |
| FASTCORE | 732 | 555 | 544 |
| minNW | 650 | 487 | 509 |
| GIMME | 546 | 1211 † | 1165 † |

\* As iGD726 contains an updated biomass with a more complex membrane lipid composition, this model is expected to have more metabolites than core models produced from iGD1575.
† The high number of reactions/metabolites is at least partially due to the presence of the complete complement of exchange reactions.

The ability of the core models to capture context-specific core metabolism was examined by predicting the essentiality of central carbon metabolic genes (Figure 3). Results were compared to both the full iGD1575 model and to the manually constructed *S. meliloti* iGD726 core metabolic reconstruction [27]. The entire set of central carbon metabolic pathways was predicted to be non-essential in iGD1575 presumably due to network redundancy. In contrast, most of central carbon metabolism was essential in the manually prepared iGD726 core model (*gnd* and *tal* are correctly predicted as non-essential). Using only Tn-seq data, Tn-Core extracted a core model largely consistent with iGD726, although the ATP synthase pump was missing. However, by also including RNA-seq data in the pipeline, the extracted core model even better reflected context-specific metabolism. This is highlighted by the lower half of the Embden-Meyerhof-Parnas pathway. In particularly, mutation of *pgk* was experimentally shown to result in a 40% growth rate decrease when grown with glucose [29]. Whereas *pgk* was essential in the first core model, deletion of *pgk* in the core model extracted using Tn-seq and RNA-seq data resulted in a growth rate decrease of 30%. Taken together, these results demonstrate the ability of Tn-Core to produce highly accurate context-specific core metabolic models, and illustrates how integrating both Tn-seq and RNA-seq data sets can lead to high precision fitness predictions.

**Figure 3.**
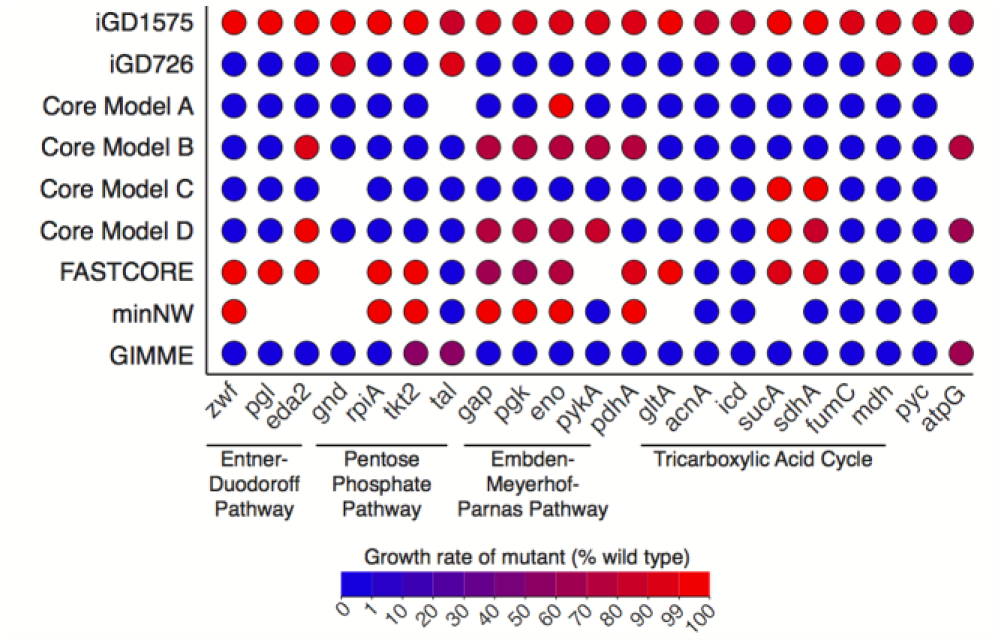
Comparison of central carbon metabolism of full and core metabolic models. This figure represents the full *S. meliloti* genome-scale metabolic reconstruction (iGD1575), the manually produced core metabolic reconstruction (iGD726), four core models produced from iGD1575 using Tn-Core (Core Model A [with Tn-seq, without RNA-seq, FBA algorithm], Core Model A [with Tn-seq, with RNA-seq, FBA algorithm], Core Model A [with Tn-seq, without RNA-seq, MOMA algorithm], Core Model A [with Tn-seq, with RNA-seq, MOMA algorithm]), and core models derived from iGD1575 using the FASTCORE, minNW, or GIMME algorithms. Representative genes from central carbon metabolism and the ATP synthase are shown. For each gene, a circle is shown if the gene is present in the model, and the circle is coloured according to the effect of deleting the gene on the growth rate of the model (determined using the MOMA algorithm); a value of 100 means no growth impact, a value of 0 means the gene deletion is lethal.

We subsequently implemented in Tn-Core the option to employ the Minimization of Metabolic Optimization (MOMA) algorithm during core model generation instead of FBA. Using MOMA instead of FBA is significantly slower, had little effect on the size of the core models (Table 1), and, at least in central carbon metabolism (Figure 3), did not produce more accurate core reconstructions. We have also found that the core models returned when using the MOMA implementation are not guaranteed to grow. This appears to be due to certain core models growing when using the *MOMA* function of the COBRA toolbox, but not growing when using the *optimizeCbModel* function of the COBRA toolbox. We therefore suggest that the FBA implementation should be used for most purposes.

The functionality of Tn-Core was further confirmed using the *Pseudomonas aeruginosa* iPae1146 GSMR [30] and published Tn-seq data [31]. These results are reported in Supplementary Material S2.

### Benchmarking of Tn-Core

There is currently no tool explicitly comparable to Tn-Core as none consider experimental Tn-seq data during core model identification. Nevertheless, we compared Tn-Core to two algorithms design for the extraction of core reconstructions: FASTCORE [10] and minNW [11]. Both algorithms are reaction-centric, and require as input a set of reactions, not genes, to be protected in the output model. To adapt these algorithms for use with Tn-seq data, we set the protected reactions as those reactions that are constrained upon deletion of the Tn-seq essential genes. Additionally, in both cases, a consistent model derived from iGD1575, generated with FASTCC [10], was used as the starting model. For both FASTCORE and minNW, the output models had similar or fewer reactions and metabolites, but a larger complement of genes, than the models produced with Tn-Core (Table 1), which is related to its reaction-centric nature. More importantly, although faster than Tn-Core, the accuracy of FASTCORE and minNW was far exceeded by Tn-Core using central carbon metabolism as a proxy (Figure 3). This result validates that Tn-Core fulfills a function that is currently lacking among the available algorithms.

The output of Tn-Core was also compared to the gene-centric TIGER implementation of the GIMME algorithm [32, 33]. GIMME generates context-specific models based on expression data, and is therefore not directly comparable to Tn-Core that primarily uses essentiality data. GIMME initially failed to return a functional model using iGD1575 and the provided RNA-seq data, but a working model could be recovered using a custom extension (see Supplementary File S2). Overall, the models returned by GIMME and Tn-Core displayed high consistency, with the central carbon metabolism extracted by GIMME of similar accuracy to those extracted by Tn-Core (Figure 3). Additionally, the GIMME model and Tn-Core model produced with Tn-seq and RNA-seq data (FBA implementation) share > 87% of their genes. Thus, at least in *S. meliloti* where essential genes tend to be highly expressed [27], both Tn-Core and GIMME perform similarly and the choice of algorithm would be driven primarily by the type of data being incorporated with the GSMR.

### Tn-Core performance

In order for Tn-Core to be accurate, a sufficiently large population of core models must be generated to ensure the optimal core model is represented. There are therefore two primary factors contributing to the speed of Tn-Core: (i) running time per iteration (i.e., per core model produced), and (ii) the number of iterations. To test the effect of starting model and parameter settings on the performance of Tn-Core, we generated 25,000 core models for five different GSMRs with varying parameter settings. A summary of these runs are provided in Table 2, and a detailed description of is reported in Supplementary File S2. 25,000 iterations did not guarantee the presence of all possible core models in any of the runs. However, the number of variably present genes gives an indication of the number of iterations required to cover all possibilities; the square of the variably present genes represents the theoretical maximum number of genetically unique core models. Considering that the variability among core models is highly dependent on the starting GSMR and the parameter settings, we recommend users first perform a test run of 10,000 iterations, and use the gene variability to approximate how many iterations must be performed. Additionally, if Tn-Core is being used to produce a core model and not only to explore redundancies in the core network, we recommend setting Tn-Core to pre-determine the essential genes prior to core model generation and to use a growth threshold of at least 50%.

**Table 2.**
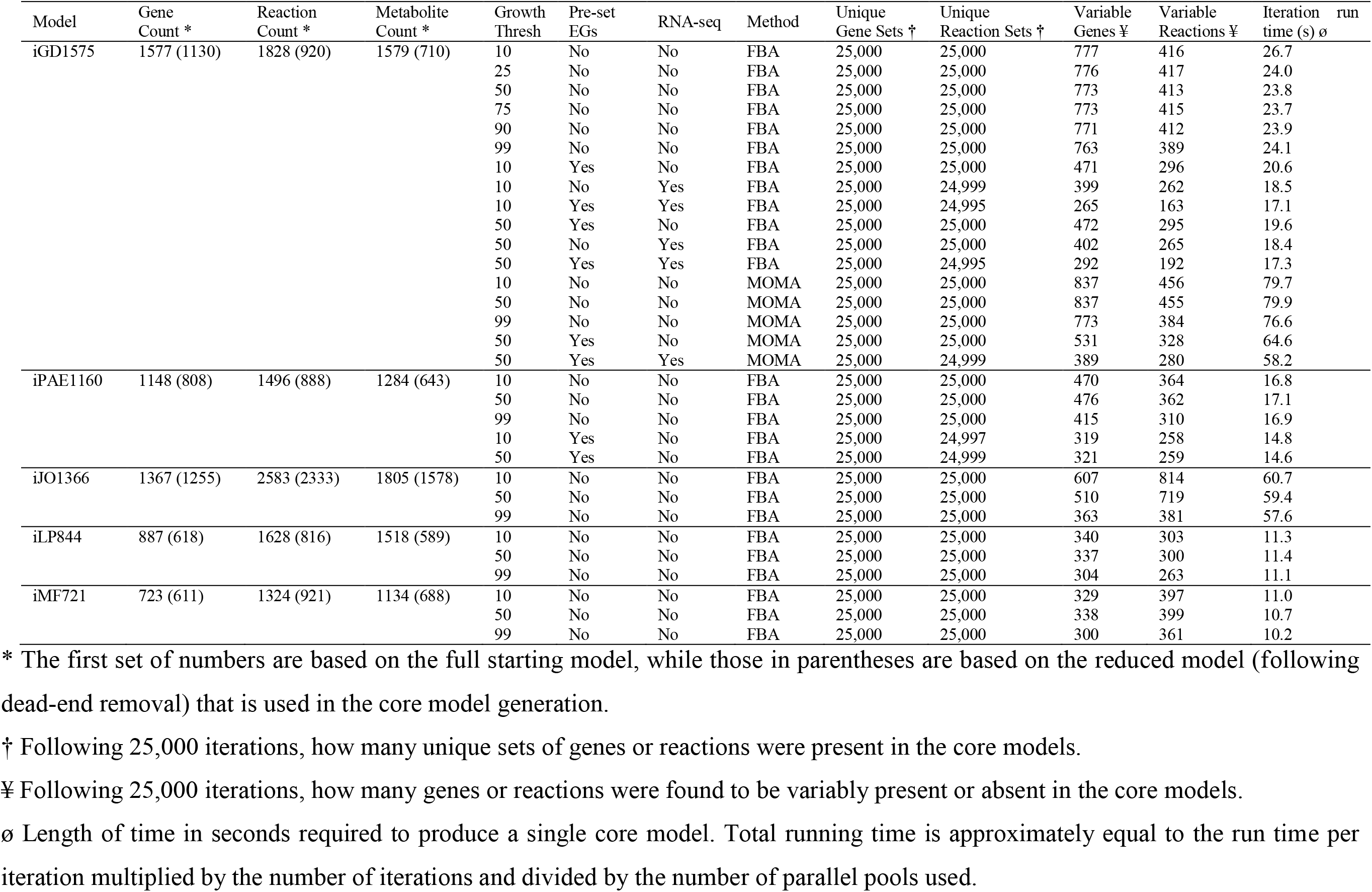
Parameters and summary statistics for Tn-Core runs.

### Characterization of redundancy and growth promoting pathways with Tn-Core

As is evident from Table 2, significant redundancy can exist in core metabolic pathways. Tn-Core produces a series of matrixes to summarize this variability (Figure 2), which can be easily imported into graphing tools to visualize the data (e.g. [34]). Here, we briefly illustrate the usefulness of these matrixes in uncovering biologically interesting data. We note that the same trends were observed for *S. meliloti* using the FBA (Figure 2) or MOMA (Figure S2) implementation, and also when using GSMRs for *Eschericha coli, P. aeruginosa, Pseudomonas haloplanktis*, and *Acinitobacter baumannii* (Figures S3-S6), demonstrating that these results are not specific to a single model (Figure S6).

Gene/reaction presence matrixes (Figures 2a, 2b) provide an overview of the variability of the models. In the case of *S. meliloti*, the core models contain an average of 434 genes, of which 286 genes (~ 66%) are invariably present and the rest are from a set of 777 variably present genes. In other words, a third of core *S. meliloti* metabolic genes can be functionally replaced by alternative genes or pathways, consistent with recent experimental work [27]. The variable and invariable core genes were mapped to KEGG pathways [35] using eggNOG-mapper [36] to identify functional biases. Significant redundancy was observed in a diversity of pathways, including carbon, amino acid and nucleotide metabolism. In contrast, the most fundamental cellular processes appeared to lack redundancy, such as transcription, translation, and aminoacyl-tRNA biosynthesis.

Gene/reaction co-occurrence matrixes summarize the frequency that two genes or reactions occur in the same model relative to chance (Figures 2c-2d). This can identify modules that work together (likely to co-occur), and genes or biochemical pathways that are functionally redundant (unlikely to co-occur). For all GSMRs used in this work, clear modules and redundant genes/pathways could be observed in the matrixes (Figure 2, Figures S2-S6). Known redundancies could be detected in the *S. meliloti* iGD1575 reaction co-occurrence matrix. For example, the two pathways for L-proline biosynthesis [37] were unlikely to occur in the same model, as were thiamine transport and thiamine biosynthesis. These observations confirm that these matrixes could be useful in detecting metabolic redundancy in core bacterial metabolism.

Finally, core models were generated using growth thresholds of 10% and 99% (of the original objective function flux), and a scatterplot was used to compare the frequency of each gene/reaction in the resulting core model populations (Figures 2f, 2g). In all cases, some genes/reactions were found to be enriched in one of the two core model populations, and the use of the MOMA algorithm increased the incidence of such genes/reactions (Figures 2 and S2). When using the FBA algorithm, biases in the occurrence of genes in the two core model populations were particularly prevalent in the *E. coli* iJO1366 model (Figure S5). Intriguingly, some genes, such as *b2417* (glucose-specific enzyme IIA component of PTS, glycolysis), *b2342* and *b3845* (both acetyl-CoA acyltransferase, fatty acid degradation), were ~ 5-fold more prevalent in the core models generated with a 99% growth threshold compared to a 10% growth threshold (differences statistically significant based on Fisher exact tests, p-value < 2.2e-16). Yet, despite the importance of the pathways these genes are involved in, none of them had a predicted effect on growth rate when deleted in the full iJO1366 model (using either FBA or MOMA), likely due to the redundancy in the complete GSMR. Hence, Tn-Core may facilitate the identification of genes contributing to optimal growth in core metabolic networks, including genes not readily detected as important in the full GSMR.

### Refinement of GSMRs using Tn-Core

Automated metabolic network reconstruction methods are expected to incorrectly assign multiple genes to the same core metabolic reaction. In the absence of experimental data, it can be difficult to correct such errors. We therefore implemented a function for using Tn-Core to assist in model refinement using Tn-seq data. We tested this pipeline using the *S. meliloti* iGD1575 model, as well as with a draft *S. meliloti* model prepared using the Kbase automated reconstruction pipeline. This process resulted in the modification of the GPRs of 60 reactions in iGD1575, with 69 genes removed from the model. Similarly, 107 GPRs (over 6% of reactions) were modified in the draft model following this process, with 57 genes deleted from the model. These results demonstrate that Tn-seq data and Tn-Core can play a valuable role in curation of metabolic models, although it certainly does not replace the need of an accurate manual curation.

## CONCLUSIONS

Here, we presented Tn-Core, a new tool for the generation of core metabolic network reconstructions. The unique feature of Tn-Core is the ability to consider experimental Tn-seq data, as well as both Tn-seq and RNA-seq data, for producing a core model that best represents the true metabolism of the cell in a given physiological condition. Despite that this pipeline may run slower than existing algorithms for the generation of core or context-specific models, Tn-Core remains advantageous due to: i) its high accuracy; ii) its ability to consider both functional genomics (Tn-seq) and transcriptomics data (RNA-seq); iii) its ease of use with little pre-processing of the data required; and iv) its gene-centric approach.

## METHODS

All data generated with Tn-Core (except for the timing of Table 2) was done using Matlab 2016a (Mathworks), the COBRA Toolbox (downloaded December 9, 2016 from the openCOBRA repository) [25], and using the Gurobi 6 solver (gurobi.com), SBMLToolbox 4.1.0 [38], and libSBML 5.11.8 [39]. All other computations were performed in Matlab 2017a using the Gurobi 7.0.2 solver, SBMLToolbox 4.1.0, libSBML 5.15.0, scripts from the COBRA Toolbox (downloaded May 12, 2017 from the openCOBRA repository), and the TIGER Toolbox v.1.2.0-beta [33]. For running minNW, the iLOG CPLEX Studio 12.7.1 solver (ibm.com) was used. Gene essentiality was determined using the *singleGeneDeletion* function and the MOMA algorithm. In order to ensure that core model generation with Tn-Core did not occasionally fail when using the MOMA algorithm, the MOMA.m script of the COBRA Toolbox was modified at line 216 to to treat unbounded solutions the same as infeasible solutions. Additionally, the solveCobraQP.m script of the COBRA Toolbox was modified to work with the Gurobi 6 solver. Detailed usage, and modifications, of FASTCORE [10], minNW [11], and GIMME [32, 33] are provided in Supplementary Materials S2.

The *S. meliloti* iGD1575 [26], *P. haloplanktis* iMF721 [40], *A. baumannii* iLP844 [41], *E. coli* iJO1366 [42], and *P. aeruginosa* iPae1146 [30] models were previously published. Prior to using iLP844, the genes ‘Unknown1’ through ‘Unknown160’ were replaced with a single gene called ‘Unknown’. The draft *S. meliloti* GSMR was generated using Kbase (kbase.us) as described in Supplementary Materials S1.

Scripts to repeat all benchmarking, as well as all output data generated in this work, are available at https://github.com/diCenzo-GC/Tn-Core. The complete Tn-Core toolbox, together with a reference manual, are provided as Supplementary Materials S1. Tn-Core is also freely available at https://github.com/diCenzo-GC/Tn-Core, and future releases of the toolbox will be available through the same link.

## ACKNOWLEDGEMENTS

GCD was funded by a Natural Sciences and Engineering Research Council (NSERC) of Canada Postdoctoral Fellowship.

